# Spatial decomposition of ultrafast ultrasound images to identify motor unit activity – A validation study using intramuscular and surface EMG

**DOI:** 10.1101/2023.06.21.545924

**Authors:** Robin Rohlén, Emma Lubel, Bruno Grandi Sgambato, Christian Antfolk, Dario Farina

**Affiliations:** Department of Biomedical Engineering, Lund University, Lund, Sweden; Department of Bioengineering, Imperial College London, London, UK

**Keywords:** ultrafast ultrasound, electromyography, motor units, spike-triggered averaging, blind source separation

## Abstract

The smallest voluntarily controlled structure of the human body is the motor unit (MU), comprised of a motoneuron and its innervated fibres. MUs have been investigated in neurophysiology research and clinical applications, primarily using electromyographic (EMG) techniques. Nonetheless, EMG (both surface and intramuscular) has a limited detection volume. A recent alternative approach to detect MUs is ultrafast ultrasound (UUS) imaging. The possibility of identifying MU activity from UUS has been shown by blind source separation (BSS) of UUS images. However, this approach has yet to be fully validated for a large population of MUs. Here we validate the BSS method on UUS images using a large population of MUs from eleven participants based on concurrent recordings of either surface or intramuscular EMG from forces up to 30% of the maximum voluntary contraction (MVC) force. We assessed the BSS method’s ability to identify MU spike trains from direct comparison with the EMG-derived spike trains as well as twitch areas and temporal profiles from comparison with the spike-triggered-averaged UUS images when using the EMG-derived spikes as triggers. We found a moderate rate of correctly identified spikes (53.0 ± 16.0%) with respect to the EMG-identified firings. However, the MU twitch areas and temporal profiles could still be identified accurately, including at 30% MVC force. These results suggest that the current BSS methods for UUS can accurately identify the location and average twitch of a large pool of MUs in UUS images, providing potential avenues for studying neuromechanics from a large cross-section of the muscle. On the other hand, more advanced methods are needed to address the non-linear summation of velocities for recovering the full spike trains.

## Introduction

The central nervous system initiates voluntary force production and human movement by delivering excitatory inputs to motoneurons that are subsequently transmitted to muscle fibres via synaptic connections. The motoneuron and innervated fibres constitute a motor unit (MU), which is the smallest voluntarily controlled structure of the human body (Heckman and Enoka, 2012). MUs have been extensively studied in basic research and clinical applications (Brown et al., 2000). Yet, the knowledge about the *spatiotemporal* neuromechanics of a large active pool of MUs in *superficial* and *deep* muscles is limited due to technological constraints.

Analysing MU activity in humans has mainly been done via analysis of electrical activity using invasive (Adrian and Bronk, 1929; Daube and Rubin, 2009) or non-invasive (Farina et al., 2014; Fuglevand et al., 1992) electromyography (EMG) techniques. These techniques mainly analyse the characteristics of the discharge times of the MU action potentials, i.e., temporal information. Although scanning-EMG, an extension of the needle EMG technique, has previously been used to study *spatial* distributions of a MU (Diószeghy, 2002; Stålberg and Antoni, 1980), the EMG techniques have limited information for a large cross-section in muscles where surface EMG is limited in depth from the skin and intramuscular EMG is limited in volume around the electrode.

Recently, ultrafast ultrasound (UUS) has been used to image and analyse individual MU activity in vivo during isometric contractions (Carbonaro et al., 2022; Lubel et al., 2022; Rohlén et al., 2020a). This technique provides sub-millimetre resolution, kHz-frame rate, and access to a large cross-section in the muscle. The approach is based on 1) calculating velocity (displacement) fields from the recorded ultrasonic signals and 2) decoding the velocity field into individual MU synchronised velocities (Rohlén et al., 2020b, 2020a). In this way, the method provides an estimate of the spatial distribution of the fibres of a MU and of the subtle mechanical oscillations in response to the fibres’ depolarisation (twitches) in both superficial and deep muscles.

The velocity field associated with individual MUs can be extracted by spike-triggered averaging (STA) of the ultrasound images using the time instants of activation of MUs identified with concurrently recorded EMG (Lubel et al., 2023, 2022). Alternatively, the velocity images associated with muscle contractions can be directly decoded into contributions of individual MUs (Rohlén et al., 2020b, 2020a). In this way, UUS can be used to estimate both the discharge times and velocity fields of individual MUs.

Direct decoding MU activity from UUS has the potential to become a new independent method for the study of the neural and mechanical characteristics of individual MUs. MU decoding from UUS has been attempted with blind source separation (BSS) (Rohlén et al., 2020b, 2020a). This approach is computationally fast and suitable for real-time processing (Rohlén et al., 2023c). However, the BSS method has yet to be validated for a large population of MUs. Previously, a study compared the BSS outputs with MU activities based on concurrent recordings with a concentric needle electrode (Rohlén et al., 2020a). Nevertheless, the mechanical coupling between the needle and the muscle tissue may have limited this validation to a very small number of MUs (Maitland et al., 2022; Rohlén et al., 2020a).

This study aimed to validate the BSS method (Rohlén et al., 2023c) to identify individual MU activity in UUS recordings using a large population of MUs. The validation was achieved by assessing its ability to identify firings, twitch areas and temporal profiles in the tibialis anterior (TA) of eleven healthy participants with concurrent recordings of either high-density surface EMG (HDsEMG) or fine-wire intramuscular EMG (iEMG). The MU population was decoded from low force levels when using HDsEMG and for a broader range of forces, up to 30% of maximum voluntary contraction (MVC) force, when using iEMG.

## Methods

### Experimental procedures

This study comprises two separate datasets of synchronised UUS, EMG, and force recordings from isometric ankle dorsiflexion contractions targeting the TA muscle. The EMG technique that was used in the first dataset (10 subjects, 26.2 ± 2.9 years) was HDsEMG, retrospectively included from a previous study (Lubel et al., 2022). The EMG technique used in the second dataset (one subject, 32 years) was iEMG.

Both studies were conducted following the Declaration of Helsinki. The two studies were approved by the Imperial College Research Ethics Committee (reference numbers: 20IC6422 and 19IC5641). After a detailed explanation of the study procedures, informed consent was obtained from all participants.

### Experimental setup dataset 1 – High-density surface EMG and MU feedback

After shaving the skin above the muscle and applying an abrasive paste, two HDsEMG electrode arrays (5×13 electrodes, 8 mm inter-electrode distance; OT Bioelettronica, Torino, Italy) were placed over the proximal and distal parts of the TA along the fibre direction, leaving a 15-mm gap between the two arrays (Lubel et al., 2022). After securing the electrodes with Tegaderm Film Dressings and self-adhesive medical bandages, the participant sat in a chair with their leg strapped into an ankle dynamometer (OT Bioelettronica, Torino, Italy) such that the foot was at an angle of 90 degrees to the leg. Finally, using a custom-built probe holder, the ultrasound probe was placed between the two EMG arrays such that the image plane was perpendicular to the fibres, resulting in a cross-sectional view of the muscle.

A real-time online EMG decomposition algorithm (Barsakcioglu et al., 2021), was trained on a 41-s-long isometric contraction at 10% of the MVC. After training the decomposition model, the experimenter inspected the decomposition output to determine if the model was properly trained and if the noise level was acceptable, leading to repeated training where necessary to ensure accurate decoding. Then, the subject was instructed to recruit and keep a single MU active for 30 s based on visual feedback while recording synchronised UUS and HDsEMG data. This procedure was repeated three times with a 60-s rest between recordings and further repeated for an increasing number of concurrently active MUs, ranging from one (minimum) to six (maximum) for each subject.

### Experimental setup dataset 2 – Intramuscular EMG and various force levels

After cleaning the skin above the muscle, an expert used a Butterfly ultrasound probe (Butterfly Network, Massachusetts, USA) to guide a fine-wire intramuscular EMG electrode (OT Bioelettronica, Torino, Italy) along the fibre length at a 45-degree angle, such that the needle was aligned with the centre of the ultrasound probe. Once the electrode was correctly positioned, the needle was removed (leaving the thin wire electrode in place), and the wires were secured to the skin with tape. The leg was secured in the ankle dynamometer, and the ultrasound probe was placed such that the image plane was as close as possible to the needle tip in a cross-sectional view.

The participant performed an MVC contraction before performing two repeats of four force-level trapezoid ramps at 5%, 10%, 20%, and 30% MVC, with 60 s of rest between each repeat. Each ramp had a 40-s plateau with 5-s recruitment and de-recruitment ramps. Once the participant reached the plateau, a 30-s UUS recording was performed. Once one set of eight ramps had been completed, the procedure was repeated after inserting a second wire electrode medially with respect to the first electrode. Finally, a third electrode was inserted laterally, and the eight ramps were repeated, resulting in a total of 24 recordings.

### Synchronised UUS, EMG, and force recordings

The signals from the HDsEMG arrays (5×13 electrodes, 8 mm inter-electrode distance; OT Bioelettronica, Torino, Italy) were recorded in monopolar derivation, amplified, sampled at 2048 Hz, A/D converted to 16 bits with 150x gain, and bandpass filtered (10-500 Hz), using a Quattrocento amplifier (OT Bioelettronica, Torino, Italy). The force data were fed through the Forza force amplifier (OT Bioelettronica, Torino, Italy) into the auxiliary port of the Quattrocento. The signals from the fine-wire EMG electrodes (OT Bioelettronica, Torino, Italy) were also recorded using a Quattrocento amplifier at 10240 Hz and bandpass filtered (10-4400 Hz).

The UUS data were recorded (L11-4v probe, 7.24 MHz centre frequency, 28.96 MHz sampling frequency) with the Vantage Research Ultrasound System (Verasonics Vantage 256, Kirkland, WA, USA) at a frame rate of 1 kHz using single-angle plane wave imaging.

All the data was aligned in post-processing based on a trigger pulse from the Verasonics system (1 µs) that was prolonged (490 µs) via an Arduino Uno microcontroller board and sampled by the Quattrocento with the EMG sampling rate (2048 Hz for the HDsEMG and 10240 Hz for the iEMG).

### EMG decomposition

#### High-density surface EMG – Blind source separation

As described in the experimental setup for the first dataset, a real-time online EMG decomposition algorithm (Barsakcioglu et al., 2021), was used to decode MU spike trains. In short, the real-time online decomposition uses a convolutive BSS and a fixed-point iteration approach to estimate a separation matrix to extract MU spike trains from the recorded HDsEMG (Negro et al., 2016).

#### Intramuscular EMG – Template matching

An automatic template-matching method in EMGLAB (McGill et al., 2005) decomposed the full length of each iEMG signal after applying a high-pass filter of 1000 Hz. An expert visually corrected the estimated spike trains for missing or double firings. Finally, the estimated MU spike trains with inter-spike interval coefficient of variation greater than 30% were discarded (Holobar et al., 2010).

### Decoding velocity images into MU twitch areas and temporal profiles

Each 30-s UUS recording resulted in 30000 frames, consisting of 357×128 pixels (∼35 mm axially and 38 mm laterally). After delay and sum beamforming the raw ultrasound data (Fig. 1A), velocity images were computed using 2D autocorrelation velocity tracking of the radio frequency signals (Loupas et al., 1995) with two consecutive frames and a 1-mm window in the axial direction (Fig. 1B). After downsampling axially and removing regions of no interest (>12 mm for HDsEMG-synchronised data and >20 mm for iEMG), there were 29998 frames, each consisting of 39×128 (HDsEMG) or 64×128 (iEMG) pixels. A negative axial velocity implies movement away from the ultrasound probe, whereas a positive axial velocity implies movement towards the probe.

**Figure 1.**
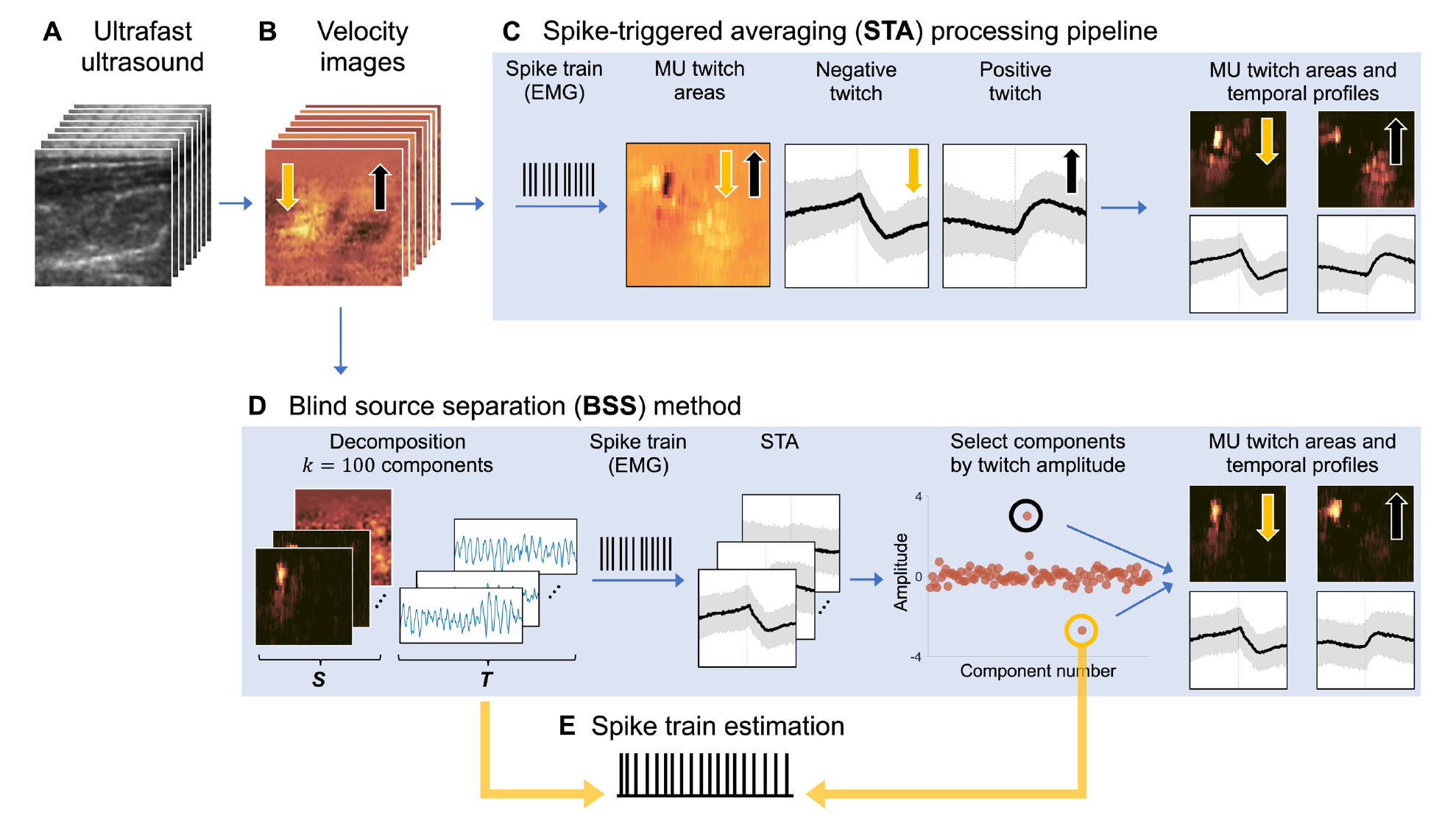
Decoding velocity images into the motor unit (MU) twitch areas and temporal profiles using the blind source separation (BSS) method. After delay and sum beamforming (**A**), 2D autocorrelation velocity tracking was performed to obtain velocity images (**B**). A negative velocity implies movement away (yellow arrow) from the ultrasound probe, whereas a positive axial velocity implies movement towards the probe (black arrow). Then, the BSS method (**D**) was applied to extract MU twitch areas and temporal profiles. As a reference, a spike-triggered averaging (STA) processing pipeline (**C**) was also used to extract MU twitch areas and temporal profiles. These were compared regarding spatial distance to the area locations and the twitches’ temporal correlation. Finally, the selected component from the BSS output was used for spike train estimation (**E**) to assess the rate of spike train agreement with spikes derived from electromyography (EMG).

#### BSS method for UUS

Each velocity image sequence was decomposed into *MU twitch areas* and *temporal profiles* using a BSS method (Fig. 1D). This decomposition was achieved using a linear mixture approach, i.e., assuming the summation of each source consisting of a (sparse) spatial map multiplied by an oscillatory temporal response (Rohlén et al., 2022b, 2020a, 2020b). Although a convolutive BSS approach, similar to the ones used on HDsEMG successfully (Holobar and Zazula, 2007; Negro et al., 2016), could be more suitable to provide the spike train directly from the velocity images, it is computationally demanding, particularly for UUS data. Also, the algorithms decoding performance of velocity images are unknown. Therefore, we used a fast spatial BSS method based on a linear model (Rohlén et al., 2023c) similar to the ones used in previous studies (Carbonaro et al., 2022; Rohlén et al., 2020a).

In short, the mathematical model can be summarised as the following. The velocity image sequence ***Y*** containing the axial velocities can be represented as a two-dimensional data matrix through vectorisation with the size of *n_s_* × *n_t_*, where *n_s_* = *n_x_* × *n_y_* is the number of pixels for the lateral and depth axis and *n_t_* is the number of frames. The decoding was based on the independent component analysis (ICA) model and a fixed-point iteration scheme under the assumption of independent and non-Gaussian spatial components (Hyvärinen et al., 2001). Here the ICA model was defined as ***Y*** = ***ST***^T^, where ***S*** is a vectorised spatial matrix and ***T*** is a temporal matrix in which a subset of the components are assumed to be *MU twitch areas* with an oscillatory signal containing a sequence of twitches (*temporal profiles*).

First, the velocity image sequence ***Y*** were z-scored along the time dimension, and the mean was subtracted in both dimensions. Second, random SVD (with exponent parameter *q* = 1 and oversampling parameter *p* = 5 (Halko et al., 2011; Rohlén et al., 2023c)) was applied to decompose ***Y*** into *k* = 100 (Rohlén et al., 2020b) left and right singular vectors. Third, ***Y*** were whitened (Hyvärinen et al., 2001) based on the left and right singular vectors. Fourth, a fixed-point iteration scheme was used to estimate a separation matrix ***w***. To estimate ***S*** (dimension *n_s_* × *k*) and ***T*** (dimension *n_t_* × *k*), we used a skewness-based gradient function *g*(*x*) = *x*^3^/3 with the first derivative equal to *g*′(*x*) = *x*^2^. Finally, the estimated ***S*** and ***T*** were obtained by projecting the separation matrix ***w*** on the whitened data matrix and the right singular vectors (Rohlén et al., 2023c).

Since the estimated ***S*** and ***T*** comprise *k* = 100 spatially independent maps and the corresponding temporal responses (Fig. 1D), we need to select which of these *k* components are associated with MU activity, as shown and discussed in previous studies (Carbonaro et al., 2022; Rohlén et al., 2023a, 2020a, 2020b). Here, we used a selection procedure using EMG-identified spikes and STA (50 ms before and after each EMG-identified spike) on each temporal response in ***T***. These spike-triggered temporal profiles were averaged to obtain a *temporal twitch profile*. Then, we calculated the STA peak-to-peak amplitudes and multiplied them with a negative sign when the direction of the motion was away from the probe after the time of the spike trigger (Lubel et al., 2022). Finally, we selected the largest *positive* and *negative* peak-to-peak STA amplitudes as the candidate *MU twitch areas* and *temporal profiles*. Fig. 1D shows an overview of this procedure.

#### STA processing pipeline

As a reference for the *MU twitch areas* and *temporal profiles* obtained from the BSS method, an STA processing pipeline (Lubel et al., 2022), was also used to decode each velocity image sequence into *MU twitch areas* and *temporal profiles* (Fig. 1C). Each spike in an EMG-identified spike train was used to extract a sequence of z-scored velocity images within a time window (50 ms before and after each spike). These spike-triggered image sequences were averaged to obtain a *spatiotemporal twitch profile*. Then, the MU twitching area was computed as the pixel-wise sum of the division between the spatiotemporal twitch profile and the variance of all the spike-triggered image sequences. The sum, i.e., a 2D map that we refer to as a *MU twitch area*, was finally multiplied with a negative sign when the direction of the motion was away from the probe after the time of the spike trigger (Lubel et al., 2022).

To extract *MU twitch profiles*, we identified the 2D pixel coordinates with the *MU twitch area*’s largest (positive) and smallest (negative) values. We found that using the ten largest/smallest values (∼1 mm^2^) empirically provided the most robust results. Then, the pixel coordinates in the *spatiotemporal twitch profiles* were averaged over all the pixel coordinates to obtain negative (away from the probe) and positive (towards the probe) *MU twitch profiles*. Fig. 1C shows these procedures.

### Accuracy of UUS BSS in spike identification

After decoding velocity images using the *BSS method* and selecting a representative component (Fig. 1D), we estimated the spike train of the selected component in the temporal matrix ***T*** (Fig. 1E) using a previously validated blind deconvolution method (Rohlén et al., 2023b). The quantification of the rate of agreement (RoA) between estimated and EMG-identified spikes was performed by using a common spike train agreement metric defined as 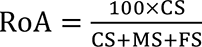, where CS was the number of correctly identified spikes, MS were the number of missed spikes, and FS the number of false spikes. The tolerance window for correctly identified spikes was set to ±15 ms (Rohlén et al., 2023b).

### Accuracy of UUS BSS in twitch area and temporal profile identification

We also assessed the MU twitch areas and temporal profiles obtained by BSS with the estimates from the EMG-derived STA processing pipeline (Fig. 1C-D). *Temporally*, we calculated the correlation between each STA-BSS-output pair of negative and positive MU twitch profiles. Complementary to the correlation, we compared the peak-to-peak amplitudes of the STA-BSS-output pairs. *Spatially*, we calculated the Euclidean distance between the median coordinates of the 2D pixels of the STA-BSS-paired *MU twitch areas* with the 10 largest and smallest values. As described in a previous section, we used ten pixels in the STA method since it empirically provided the most robust results.

### Statistical analysis

Descriptive statistics of the rate of spike train agreement, temporal correlations and spatial distance were quantified using the mean, standard deviations, minimum, and maximum values. An ordinary least squares model was used to test the linear relation (*R*^2^ and *p*-value) between 1) the peak-to-peak twitch amplitudes from the BSS method and the RoA values and 2) the peak-to-peak twitch amplitude values from the BSS method and the STA processing pipeline (both negative and positive twitches). A *p*-value of 0.05 was set for statistical significance.

## Results

After removing recordings based on poor quality, synchronisation, and system issues, 319 MUs were found from HDsEMG signals, and 83 MUs were found from iEMG signals, i.e., 402 MUs in total. By decomposing the UUS data using the BSS method, selecting the representative components, and estimating the spike trains, MU twitch areas, and temporal profiles, we found a weak spike train agreement (34.0 ± 15.0%), with 87% being the highest (Fig. 2). We found that the number of correctly identified spikes (with respect to the number of EMG-identified firings) was 53.0 ± 16.0%, the number of missed spikes was 47.0 ± 16.0%, and the number of falsely identified spikes was 58.0 ± 25.0% (Fig. 2).

**Figure 2.**
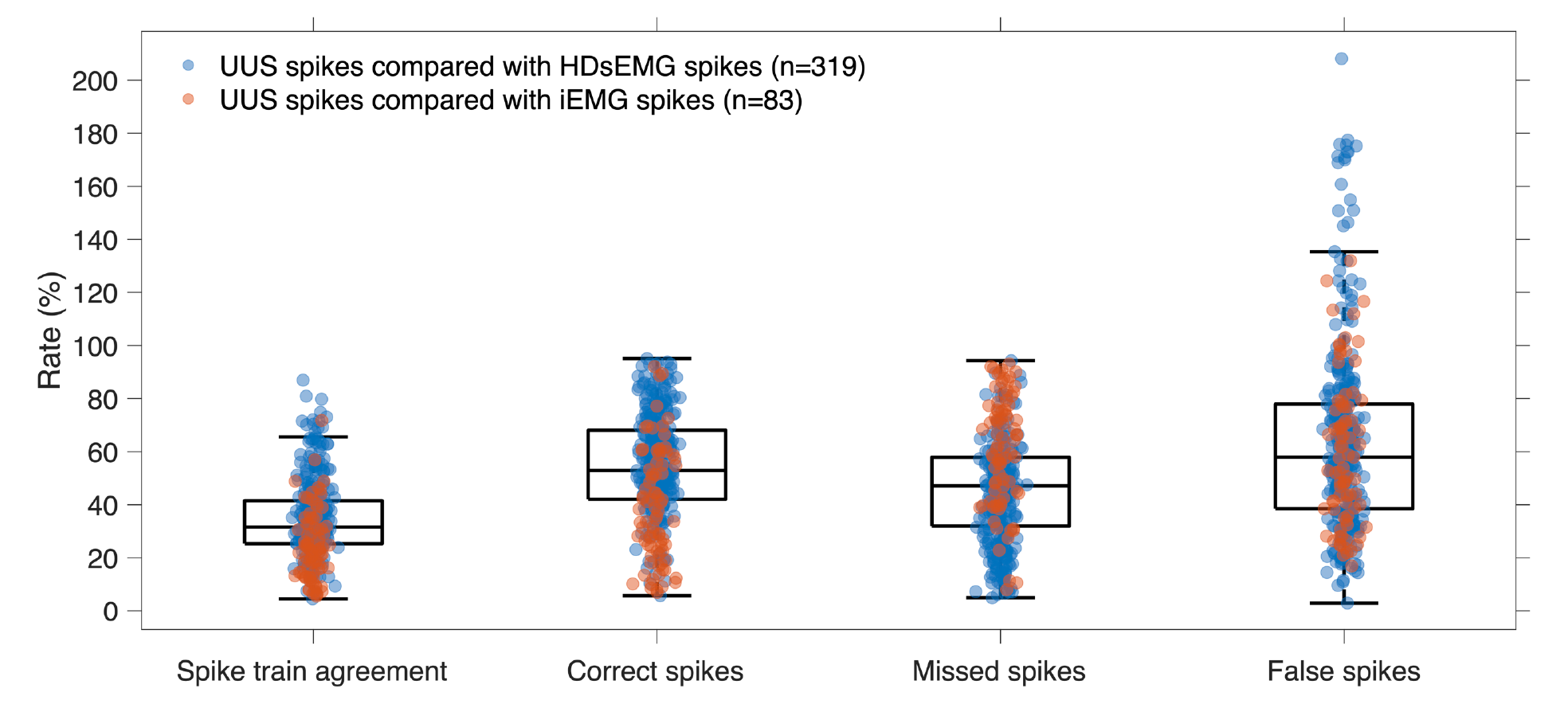
Assessment of the accuracy of identifying neural spikes using the BSS method. There was a weak spike train agreement (34.0 ± 15.0%), with the spikes from either high-density surface electrography (HDsEMG) or intramuscular EMG (iEMG), with 87% being the highest. We found that the number of correctly identified spikes (with respect to the number of EMG-identified firings) was 53.0 ± 16.0%, the number of missed spikes was 47.0 ± 16.0%, and the number of falsely identified spikes was 58.0 ± 25.0%.

The MU twitch areas from the BSS method were proximally located in space to the areas of the STA processing pipeline for both negative (1.90 ± 2.55 mm, n = 402) and positive twitches (1.84 ± 2.74 mm, n = 254 MUs). Also, the MU twitch profiles from the BSS and STA methods were highly correlated for the negative (0.94 ± 0.06) and positive twitches (0.93 ± 0.08). When analysed jointly, the space-time features of the BSS-STA pairs occupied the high correlation and low distance corners in a 2D space (Fig. 3A-B). In addition, there was a linear relation between the negative (R^2^ = 0.69; *p* < 0.001) and positive (R^2^ = 0.67; *p* < 0.001) peak-to-peak twitch amplitudes from the MU twitch profiles of BSS and STA (Fig. 3C-D).

**Figure 3.**
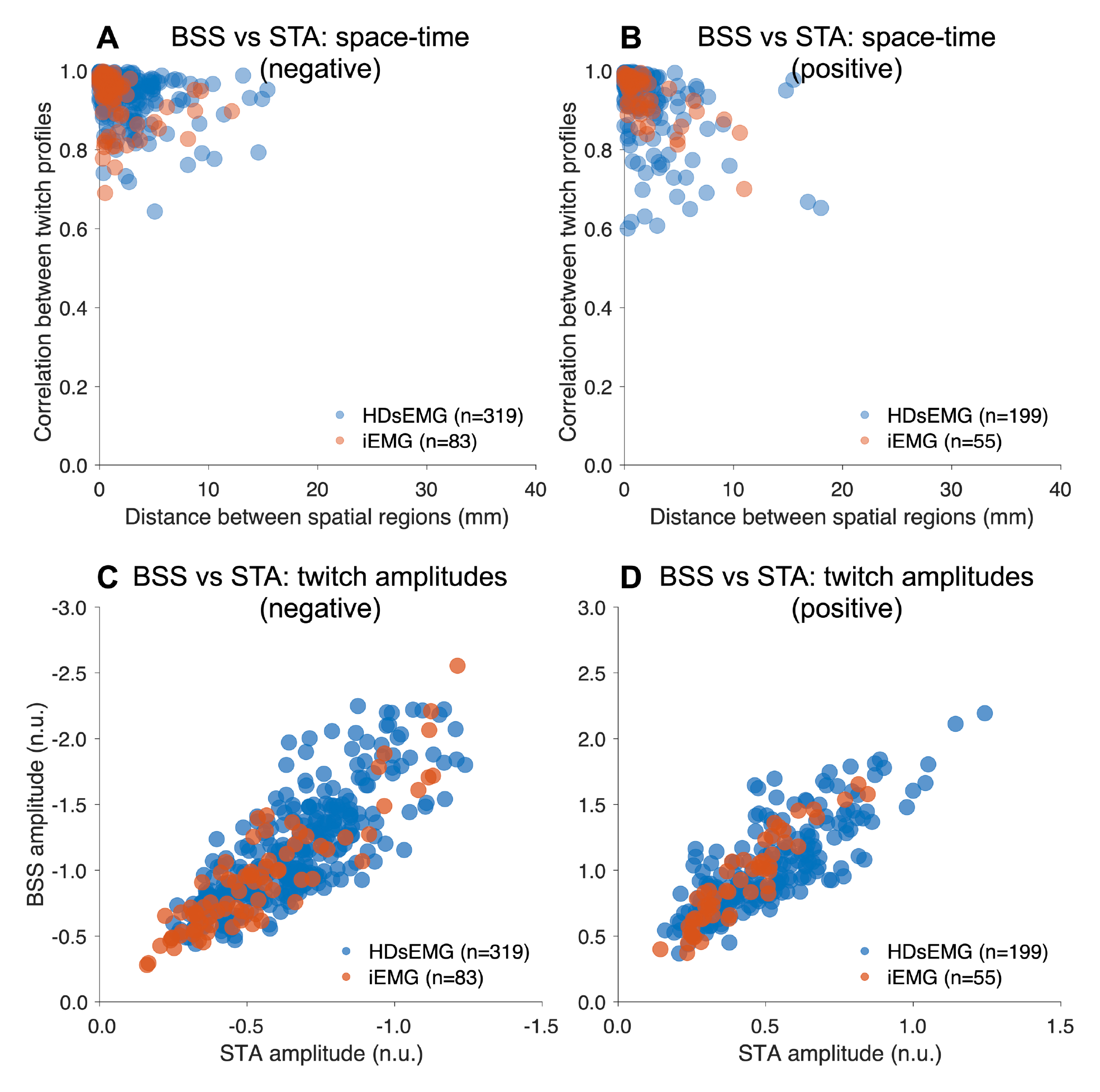
The motor unit (MU) twitch areas from the blind source separation (BSS) were proximally located in space to the ones from the spike-triggered averaging (STA) processing pipeline concerning both (**A**) negative (1.90 ± 2.55 mm) and (**B**) positive twitches (1.84 ± 2.74 mm). Also, the MU twitch profiles were highly correlated for the (**A**) negative (0.94 ± 0.06) and (**B**) positive twitches (0.93 ± 0.08). By comparing the space-time features of the BSS-STA pairs jointly, they occupy the high correlation and low distance corners in 2D space. There was a linear relationship between the (**C**) negative (R^2^ = 0.69; *p* < 0.001) and (**D**) positive (R^2^ = 0.67; *p* < 0.001) peak-to-peak twitch amplitudes from the MU twitch profiles of BSS and STA methods. There were 402 (positive) and 254 (negative) MU twitch profiles in total, considering both the high-density surface electromyography (HDsEMG) and fine-wire intramuscular EMG (iEMG).

There were two representative cases found in this study. First, there were both negative and positive twitch pairs for both BSS and STA (Fig. 4). That means two proximal areas are moving in opposite directions, i.e., away and towards the probe (skin). These have been found in other studies and are called “MU twisting” (Deffieux et al., 2008; Lubel et al., 2022). Second, there was only a negative twitch for both BSS and STA (Fig. 5). A negative twitch implies that the MU twitch area moves away from the probe (skin).

**Figure 4.**
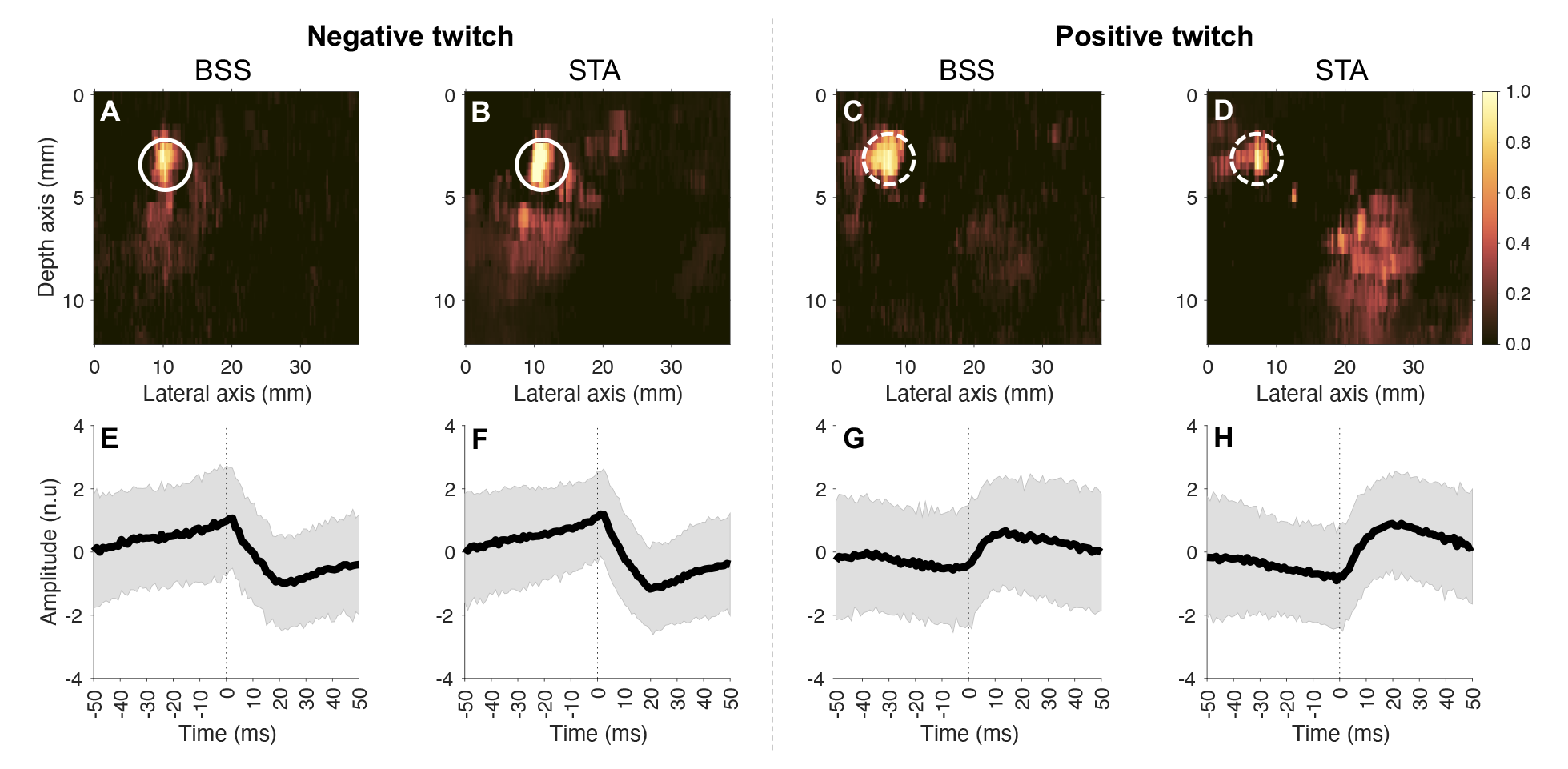
A typical example with so-called “motor unit (MU) twisting”, i.e., MU twitch areas (**A**-**D**) having negative and positive MU twitch profiles (**E**-**H**) moving away from/towards the probe (skin). The blind source separation (BSS) and spike-triggered averaging (STA) processing pipelines clearly show consistent outputs.

**Figure 5.**
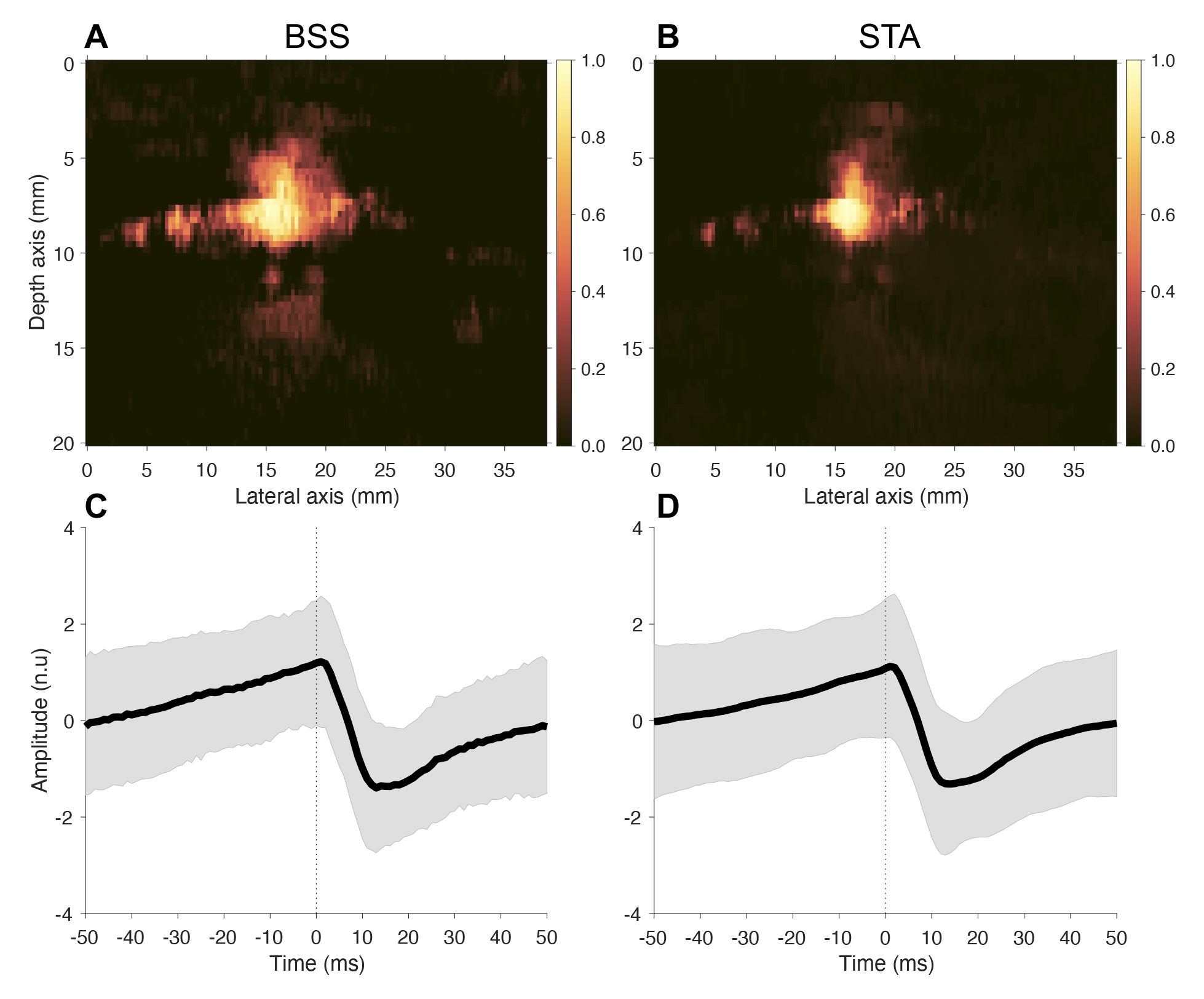
A typical example with MU twitch areas (**A**-**B**) having only negative MU twitch profiles (**C**-**D**) moving away from the probe (skin). The blind source separation (BSS) and spike-triggered averaging (STA) methods clearly show consistent outputs.

Interestingly, 10 MUs were detected at 30% MVC, where the BSS-STA pairs also were proximally located in space to each other (2.56 ± 2.45 mm), where the size of a TA and MU cross-section in healthy subjects is approximately 260 mm (McCreesh and Egan, 2011) and 5 mm (Stålberg and Dioszeghy, 1991), respectively. Also, the negative MU twitch profiles from the BSS and STA methods were highly correlated (0.89 ± 0.08, n = 10). In Fig. 6, we illustrate the consistency of the BSS and STA methods in identifying the same MU at 30% MVC in two consecutive recordings (Fig. 6).

**Figure 6.**
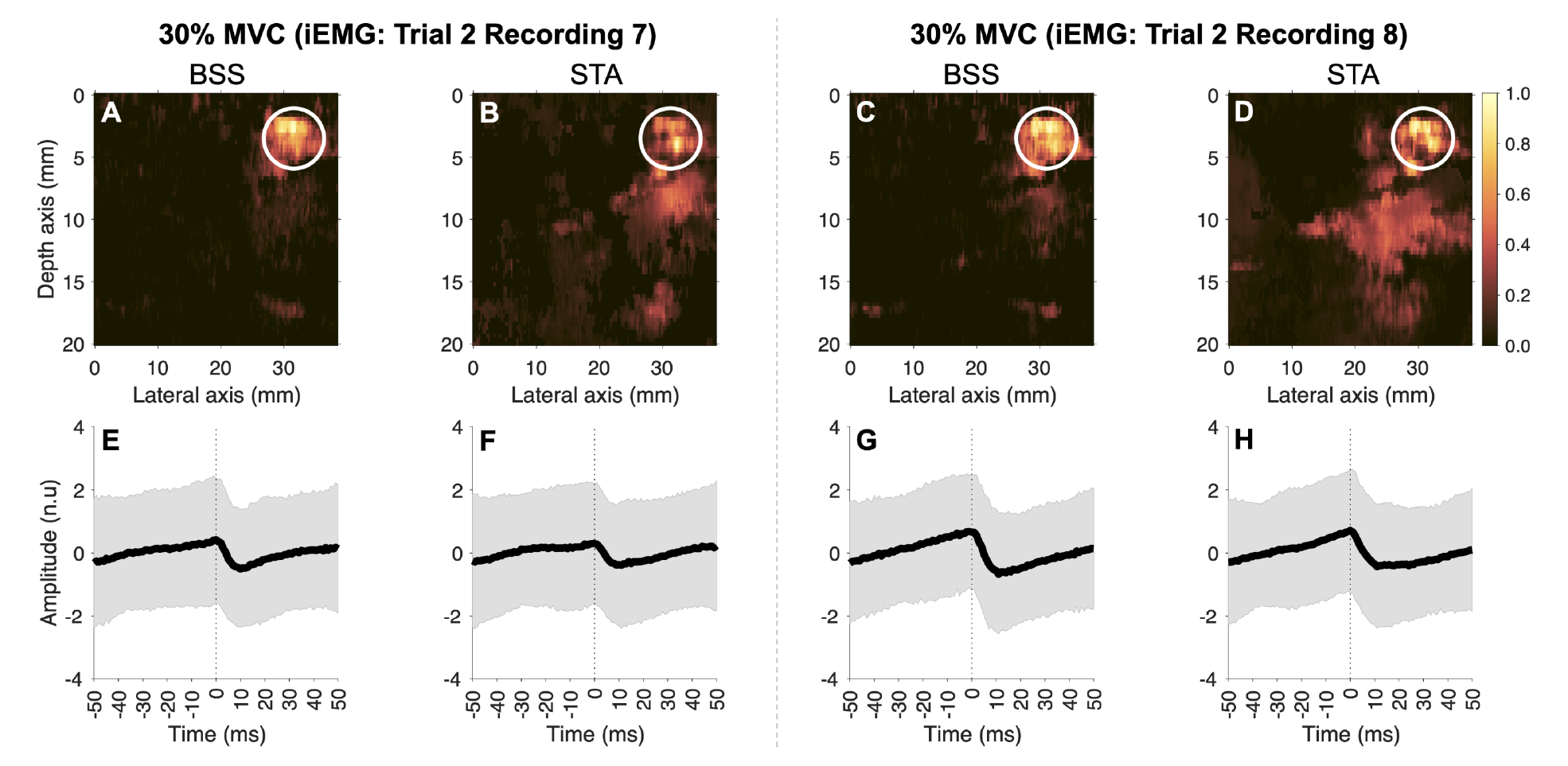
An illustration of the (**A-B**, **E-F**) blind source separation (BSS) and (**C-D**, **G-H**) spike-triggered averaging (STA) methods that can identify motor unit (MU) twitch areas and temporal profiles at 30% maximal voluntary contraction (MVC) in two consecutive recordings in the same trial. These examples were from the fine-wire intramuscular electromyography (iEMG) dataset.

## Discussion

This study aimed to validate the BSS method to identify MU spike trains, twitch areas and temporal profiles in UUS images using 402 MUs from low forces up to 30% MVC using HDsEMG or iEMG as reference methods of MU identification. We estimated spike trains using the BSS method and quantified the RoA with the detected EMG spikes. In addition, we obtained twitch areas, temporal profiles, and peak-to-peak twitch amplitudes. These spike trains, twitch areas and temporal profiles were assessed, resulting in three main findings.

First, the rate of spike train agreement with the EMG-identified firings was weak, where 53% of the firings could be identified using BSS on the UUS images (Fig. 2). The reason for this relatively low accuracy may partly be related to the non-linear summation of MU twitch velocities (displacements) in UUS, as recently shown (Lubel et al., 2023). Specifically, Lubel et al. (2023) observed that the peak-to-peak STA twitch amplitude was suppressed with increasing MU activity (force level), although the twitch waveform shape was maintained. In this study, we also found a high rate of falsely identified spikes (58%), which can be explained by additional oscillatory peaks, not in response to the spikes of the reference MU but possibly due to nearby MUs and high-amplitude noise. Hence, this study confirms that a linear mixture-based BSS model is not appropriate to identify the full neural spike train. To fully recover all the spikes of a MU, addressing the non-linear summation of velocities seems crucial.

Second, the BSS-based MU twitch areas and temporal profiles were closely located in space and had highly correlated twitches with the ones from the STA processing pipeline. The location proximity was based on selecting ten coordinates and taking the median of the minimum distances. This metric was robust for various MU twitching areas, including split territories (Birkbeck et al., 2020). We did not compare spatial distributions quantitatively because of several challenges. First, STA provides one MU twitch area (spatial map) that includes the parts of the MU that move towards the skin (positive twitch) and the parts that move away from the skin (negative twitch). In contrast, BSS may provide two separate MU twitch areas (spatial maps), one for the positive and the other for the negative twitch. In addition, BSS may divide large split regions into multiple separate components, whereas STA provides only one map. Therefore, to compare the spatial distributions, one would need additional post-processing (e.g., merging, segmenting, and mapping of the areas), subject to various parameter selections. Although comparing spatial distributions would be useful, there was a very small deviation in spatial locations and a very high correlation between the twitch profiles. This suggests that the BSS method can accurately extract MU twitch areas and temporal profiles.

Third, the BSS method can identify MUs at 30% MVC. This is the first study showing that MUs at 30% MVC can be identified using UUS, as previous studies have been constrained to very low force levels with single MU feedback (Lubel et al., 2022; Rohlén et al., 2020a) and from 2% up to 20% MVC (Carbonaro et al., 2022; Lubel et al., 2023). We expect declining performance at the higher force levels due to inherent non-linearities complicating the velocity field (Lubel et al., 2023) and violating the linear mixture assumption in both the BSS and STA methods. Going from a linear to a nonlinear BSS model (Hyvärinen et al., 2023b, 2023a) may possibly overcome this limitation by integrating prior velocity field information and, as discussed above, it is also important to estimate all the spikes accurately.

Although the BSS method relies on EMG as a reference technique to select relevant decomposed components from the BSS output, it has the potential to be a stand-alone method. For example, a classifier could be trained to classify components into MU- or non-MU- components. One may also consider replacing the EMG spike train with an estimated one using the BSS output (Rohlén et al., 2023b, 2022a) and use prior information, such as the spatial autocorrelation (as clearly visible in, e.g., Fig. 4, 5, and 6) and the typical shape of a twitch. Examples of non-MU components (which lack spatial autocorrelation) can be found in previous studies (Rohlén et al., 2023c, 2023b). Stand-alone approaches were suggested in previous studies (Ali et al., 2020; Rohlén et al., 2020b). However, while they provided good results when applied to simple linear-based simulation modelling signals, the translation to real data resulted in poor performance.

In conclusion, this study aimed to validate the BSS method to estimate individual MU activity in UUS images using 402 MUs obtained from HDsEMG and iEMG. The results of this study suggest that the current BSS method can accurately identify the location and average twitch of a large pool of MUs in UUS images, providing potential avenues for studying neuromechanics from a large cross-section of the muscle. However, to fully recover all spikes of a MU, addressing the non-linear summation of velocities is crucial.

### Competing interests

The authors declare that they have no financial and non-financial competing interests.

## Acknowledgements

The work was supported by funding from the European Union’s Horizon 2020 research and innovation programme under Grant Agreement No.899822, SOMA project. R.R. is supported by the Hans Werthén Foundation and the Swedish Research Council for Sport Science (D2023- 0003).

## Notes

### Competing Interest Statement

The authors have declared no competing interest.

